# Assessing anatomical changes in male reproductive organs in response to larval crowding using micro-computed tomography imaging

**DOI:** 10.1101/2021.07.22.453343

**Authors:** Juliano Morimoto, Renan Barcellos, Todd A. Schoborg, Liebert Parreiras Nogueira, Marcos Vinicius Colaço

## Abstract

Ecological conditions shape (adaptive) responses at the molecular, anatomical, and behavioural levels. Understanding these responses is key to predict the outcomes of intra- and inter-specific competitions and the evolutionary trajectory of populations. Recent technological advances have enabled large-scale molecular (e.g., RNAseq) and behavioural (e.g., computer vision) studies, but the study of anatomical responses to ecological conditions has lagged behind. Here, we highlight the role of X-Ray micro-computed tomography (micro-CT) in generating *in vivo* and *ex vivo* 3D imaging of anatomical structures, which can enable insights into adaptive anatomical responses to ecological environments. To demonstrate the application of this method, we manipulated the larval density of *Drosophila melanogaster* flies and applied micro-CT to investigate the anatomical responses of the male reproductive organs to varying intra-specific competition levels during development. Our data is suggestive of two classes of anatomical responses which broadly agree with sexual selection theory: increasing larval density led to testes and ejaculatory duct to be overall larger (in volume), while the volume of accessory glands and, to a lesser extent, ejaculatory duct decreased. These two distinct classes of anatomical responses might reflect shared developmental regulation of the structures of the male reproductive system. Overall, we show that micro-CT can be an important tool to advance the study of anatomical (adaptive) responses to ecological environments.

## Introduction

Animals respond to their environment across all levels of biological organisation, from gene expression through anatomical changes to complex behaviours. Recent technological advances have enabled large-scale studies of molecular and behavioural responses to ecological conditions. For instance, the advent of the RNAseq technique has provided insights into how organisms respond to key ecological conditions, including temperature (Smith et al. 2013; Kumar et al. 2020), pesticides (Christen et al. 2018; Colgan et al. 2019), social status (Veiner et al. 2021), and diet (Xu et al. 2018). Likewise, the advent of computer visions and automated tracking alrogithms has enabled studies that identify key behavioural responses in controlled and natural conditions of individuals and groups (Ferre et al. 2009; Weinstein 2018; Shreesha et al. 2020 anatomical, physiological, and developmental ;Lürig et al. 2021), allowing for a deeper understanding of behavioural responses across ecological environments. To date, however, it remains challenging to conduct anatomical studies of internal organs *in vivo* and *ex vivo*, and to some extent, our understanding of anatomical responses to ecological environments has lagged behind. Yet, understanding anatomical responses to ecological conditions can aid our understanding and predicitons of functional responses that shape the evolution of populations (Minelli 2003). It is therefore essential that new technologies are used in the field of ecology and evolution which enables the study of anatomical responses to ecological conditions.

Microcomputed Tomography (Micro-CT) is an X-ray based imaging technique that allows for non-destructive, three dimensional reconstructions of small objects (microns to millimetres) in their native states (Lin et al. 2019). This is largely due to the relatively high power of X-Rays and their ability to penetrate objects of much higher densities than visible or infrared light waves that are used in traditional imaging methods, such as light microscopy. A 3D image is obtained by rotating the object and acquiring a series of two dimensional projection images, which are then reconstructed into tomograms containing 3D information that is isotropic in its resolution using specialized algorithms (Feldkamp et al. 1984). As a result, this technique has proven useful in a number of different scientific disciplines, including biology, engineering, physics, materials science, geology, and anthropology, where dissections or destruction of an object that would be required for light imaging is not feasible (Carlson 2006; Metscher 2009a; Schambach et al. 2010; Abel et al. 2011; Rawson et al. 2020). In the biological sciences, micro-CT has proven especially useful for the nondestructuve imaging of insects, given their size and the inability of light waves to penetrate through the often thick and pigmented cuticle. Importantly, the information derived from micro-CT tomograms has proven especially useful for answering a diverse range of questions across the insect biological spectrum, from biomedical related applications utilizing highly characterized model insects such as (but not limited to) *Drosophila* (Mattei et al. 2015; Chen et al. 2018; Schoborg et al. 2019; Schoborg 2020; Khezri et al. 2021) to anatomical, physiological, and developmental applications across the wider taxonomy of insects (Westneat et al. 2003; Smith et al. 2016, 2020; Taylor et al. 2016; Chaturvedi et al. 2019; Rix et al. 2021; Wyber et al. 2021). For instance, micro-CT analysis has shown for the first time the changes in female reproductive tract following mating in *Drosophila melanogaster* (Mattei et al. 2015). Moreover, micro-CT has also been key to unravel the mechanisms of localised tissue damage in traumatic insemination in beetles (Dougherty and Simmons 2017), sperm transfer mechanism in spiders (Rix et al. 2021), and to gain insights into the evolution of genitalia morphology in lepidopterans (McNamara et al. 2019). Micro-CT has also provided the fundamental technique to assemble brain atlases of model and non-model organisms such as for example flies, bees and moths [see (Adden et al. 2020; Rother et al. 2021) and references therein]. Moreover, micro-CT has been used to reveal the anatomical and subsequent functional damage caused by pesticide exposure at different life stages in bumblebees, whereby exposure to pesticides not only reduced mushroom body calycal growth but also lower condition-learning and response to rewarding stimuli (Smith et al. 2020). Therefore, micro-CT has incredible potential to assist the morphological analysis of adaptation to changing ecological conditions (Dougherty and Simmons 2018). However, despite Micro-CT’s usefulness across a range of biological disciplines, ecological applications have remained underutilized despite the wealth of ecological knowledge that could be derived from these studies (Gutiérrez et al. 2018).

In this study, we add to the growing use of micro-CT in entomology and demonstrate how 3D micro-CT imaging can be applied to the study of anatomical responses to ecological conditions in male *Drosophila melanogaster* as a model. We demonstrate the application of micro-CT in a study-case, where we investigated the anatomical responses of the male reproductive system in the *Drosophila melanogaster* model to increasing levels of intra-specific competition at the developmental stage (i.e., number of larvae per gram of food, henceforth referred to as ‘larval density’). In holometabolous insects such as *D. melanogsater*, larval density is an important ecological factor shaping individual fitness and can be an important ecological cue of intraspecific competition levels which individuals – particularly males – are likely to encounter in the adult stage (Johnson et al. 2017). In *Drosophila*, males from high larval densities are known to have smaller body sizes and lower mating and reproductive success (Amitin and Pitnick 2007; Morimoto et al. 2016, 2017) but to have disproportionately higher ejaculate investment relative to body size in each mating opportunity (Wigby et al. 2015), potentially as a mitigatory (adaptive) response to the morphological constraints imposed by nutrient limitation and competition at the larval stage (Klepsatel et al. 2018). This effect appears to be above and beyond changes in sperm length or transcriptional levels of nine major seminal fluid proteins (Amitin and Pitnick 2007; McGraw et al. 2007). Therefore, it is possible that male reproductive responses to larval intraspecific competition originate in changes in male’s morphology (rather than physiology) that enables males to invest relatively more ejaculate per mating opportunity.

In other insects, anatomical changes such as increased testes sizes are known responses to increased larval density [e.g., (Gage 1995; Stockley and Seal 2001)], but little is known about other aspects of the male reproductive system. Moreover, non-sperm components of the ejaculate (e.g., seminal fluid proteins) which are produced in tissues other than testes can be as important for male fertility (Perry et al. 2013), but for which little is known as to how these organs respond to males’ developmental environment. In this study, we manipulated larval density within the range observed in our previous ecological assessment of *D. melanogaster* larvae in a natural population (Morimoto and Pietras 2020) and included densities ranging from low, natural range, and high larval densities relative to the densities observed in nature. This approach allowed us to apply our 3D micro-CT technique to an ecologically relevant experimental design for the species. Overall, sexual selection theory generates the implicit theoretical prediction that as larval density increases, males should allocate more (of the fewer) resources per capita to traits related to either migration (e.g., longer wings) or (post-copulatory) reproductive success (e.g., better ejaculates), or both (Katsuki et al. 2013). Thus, we predicted that male reproductive organs’ volume – particularly accessory glands and testes – should be positively associated with larval density assuming that (1) relatively larger reproductive organs are correlated with higher ejaculate investment, (2) larval density is an ecological cue for the level of (post-mating) competition in the adult stage and (3) males respond adaptively to this ecological cue as to enhance their reproductive success. Empirical work suggests that many insect species conform at least partly to this expectation [reviewed in (Than et al. 2020)]. These predictions emerged from the rationale that, as larval density increased, the perceived level of post-copulatory competition would likewise increase, resulting in high demand for high-quality ejaculates that in turn, require larger reproductive organs to produce it (Parker 2016). Overall, this study highlights the potential benefits of using micro-CT imaging as a tool to study anatomical responses to ecological conditions, helping shed light of how organisms respond to their environment and thereby evolve the myriad of forms and functions observed in the animal kingdom.

## Material and Methods

### Fly stock and larval density manipulation

We used an outbred *D. melanogaster* population collected in September 2015 in Brittany (France) and kindly provided to us by Herve Colinet (Henry et al. 2018). Flies were maintained in large population cages (>1,000 individuals) with overlapping generations, at 20 °C and ca. 50 % humidity, with 12 h light: 12 h dark cycles. Fly stocks were maintained – and all experiments conducted with a standard yeast-sucrose diet (Brewer’s yeast MP Biomedicals 0290331205, commercial sucrose, agar Sigma-Aldrich A1296, Sigma-Aldrich, and 0.5 % Nipagin Sigma-Aldrich). Eggs were collected for 6 h using an oviposition device (Petri dish (90 mm) covered with a solution of commercial blackcurrant juice and 1 % agar and coated with a thin layer of yeast paste). Oviposition devices were incubated overnight until eggs hatched at 25 °C, after which, first instar larvae were counted and allocated to larval density treatments using a soft brush under a Leica M9i stereoscope. Larval density treatments were based on our survey of larval densities in a natural population of *D. melanogaster* (Morimoto and Pietras 2020). We had 5 larval densities: 0.5, 5, 15, 30, and 50 larvae/g of diet in vials with between 3 g (highest density) to 6 g (lowest density) of diet. This larval density gradient corresponds to a range from low (i.e., 0.5 larvae/g, lower than the natural range), natural range (5 and 15 larvae/g), and high (30 and 50 larvae/g, higher than natural range) larval densities (Morimoto and Pietras 2020). Vials were incubated at 25 °C with 12 h light: 12 h dark cycles until adult emergence. Within 8 hours of emergence, females were discarded, males were transferred to fresh vials with the standard diet and incubated for 5 days before being killed at -20 °C. This ensured that even in the unlikely case that males mated, males had enough time to replenish their reproductive organs prior to imaging. Males were transferred to an increasing gradient of ethanol (from 40 % to 100 %, 2h in each solution) for fixation before imaging.

### Imaging and data analysis

Individual males were submerged in the iodine-ethanol (I2E) 1 % in 100 % ethanol (EtOH) solution and stored overnight, after which individuals were rinsed in 100 % EtOH three times for 10 min to wash off excess iodine. The specimens were then placed in heat-sealed pipette tips with 100 % EtOH and scanned using a SkyScan 2211 (Multiscale X-ray Nano-CT System, Bruker micro-CT, Kontich, Belgium) at Oral Research Laboratory, University of Oslo. Scanning parameters were as follows: 55 kV, 260 μA, 650 ms exposure time per projection and without the use of a physical filter. Males were scanned over 360° with rotation steps of 0.31°, resulting in 1162 projections. Each projection was averaged by 3 frames, leading to a total scan duration of about 40 min for each sample and a final voxel size of 1.40 μm. The reconstruction process for each sample was performed using the system-provided software NRecon (version 1.7.4.6). Image segmentation was done using Dragonfly software ORS (Object Research Systems Inc, Montreal, Canada, 2019; software available at http://www.theobjects.com/dragonfly). Each male reproductive organ, namely, accessory glands, testis, ejaculatory duct and bulb, were segmented individually employing semiautomatic processes and correcting for inconsistencies in the three axes to achieve optimal and accurate segmentation.

### Statistical analyses

All statistical analyses were conducted in R (Core Team 2010). We imaged 24 randomly selected males (0.5 larvae/g: *N =* 2, 5 larvae/g: *N =* 5, 15 larvae/g: *N =* 5, 30 larvae/g: *N =* 7, 50 larvae/g: *N =* 5). Accessory glands and testes were measured individually. We compared the volumes (in *µ*m^3^) of each organ with a linear mixed model using the ‘lme4’ and ‘lmerTest’ packages (Bates et al. 2007; Kuznetsova et al. 2017) with models that included individual ID or in the case of accessory glands and testes, side (left or right) nested within individual ID as a random effect and the fixed effect larval density (fitted as a factor with 5 levels) (Table S1). P-values were obtained from the ‘anova’ function of the ‘lmerTest’ package. Post-hoc tests in linear mixed models were performed using the ‘emmeans’ package (Lenth and Lenth 2018). In *Drosophila*, larval density influences male body size (Amitin and Pitnick 2007). We therefore also included normalized abdominal volume as a fixed effect in all models to control the allometric relationship between larval density, body size, and reproductive organs, even though abdominal volume was not statistically different between larval densities (Density: F_4,17_ = 2.955, p = 0.051; Table S1). All data plots were made using the ‘ggplot2’ package (Wickham 2016).

## Results

We first identified the target organs for segmentation. Fig 1a shows an example of an image slice of a specimen used for segmentation and Fig 1b shows a 3D model reconstruction of the segmented abdomen highlighting the male reproductive system. A video with an example of the segmentation slice per slice is presented in the supplementary material (Video S1). This approach allowed to then test the effect of larval density on male reproductive organs. After controlling for abdominal volume, testes volume was statistically significantly influenced by larval density (Density: F_4, 40_ = 10.085, p < 0.001). For instance, males from the 30 larvae/g treatment had the testes with the highest volume while males from the 15 larvae/g treatment had the testes with the smallest volume. Males from the 5 larvae/g treatment had testes with intermediate volume which was not different from males from the lowest (0.5 larvae/g) or highest (50 larvae/g) larval densities (Table S1, Fig 2a). Accessory gland volume progressively decreased with increasing larval densities, although this effect was not statistically significant (Density: F_4,39_ = 2.4247, p = 0.064). Nevertheless, this trend suggested that larger accessory glands (by volume) were observed in males that emerged from the lowest and natural larval densities of 0.5, 5 and 15 larvae/g, whereas relatively smaller accessory glands were observed from males that emerged from high larval densities of 30 and 50 larvae/g (Fig 2b, Table S1). Ejaculatory bulb displayed similar trend as that of accessory glands but without reaching statistical significance (*Ejaculatory bulb:* F_4,14.93_ = 1.584, p = 0.130). Ejaculatory ducts volumes were statistically influenced by larval density (*Ejaculatory duct:* F_4,18_ = 8.462, p = <0.001). Interestingly, the ejaculatory duct followed a similar (although less accentuated) trend to that observed for the testes (Table S1, Fig 2c). Conversely, the ejaculatory bulb followed a similar trend observed in the accessory glands, whereby males emerging from high larval densities of 30 and 50 larvae/g had smaller ejaculatory bulbs by volume compared with males from other larval density treatments (Fig 2d).

**Fig 1.**
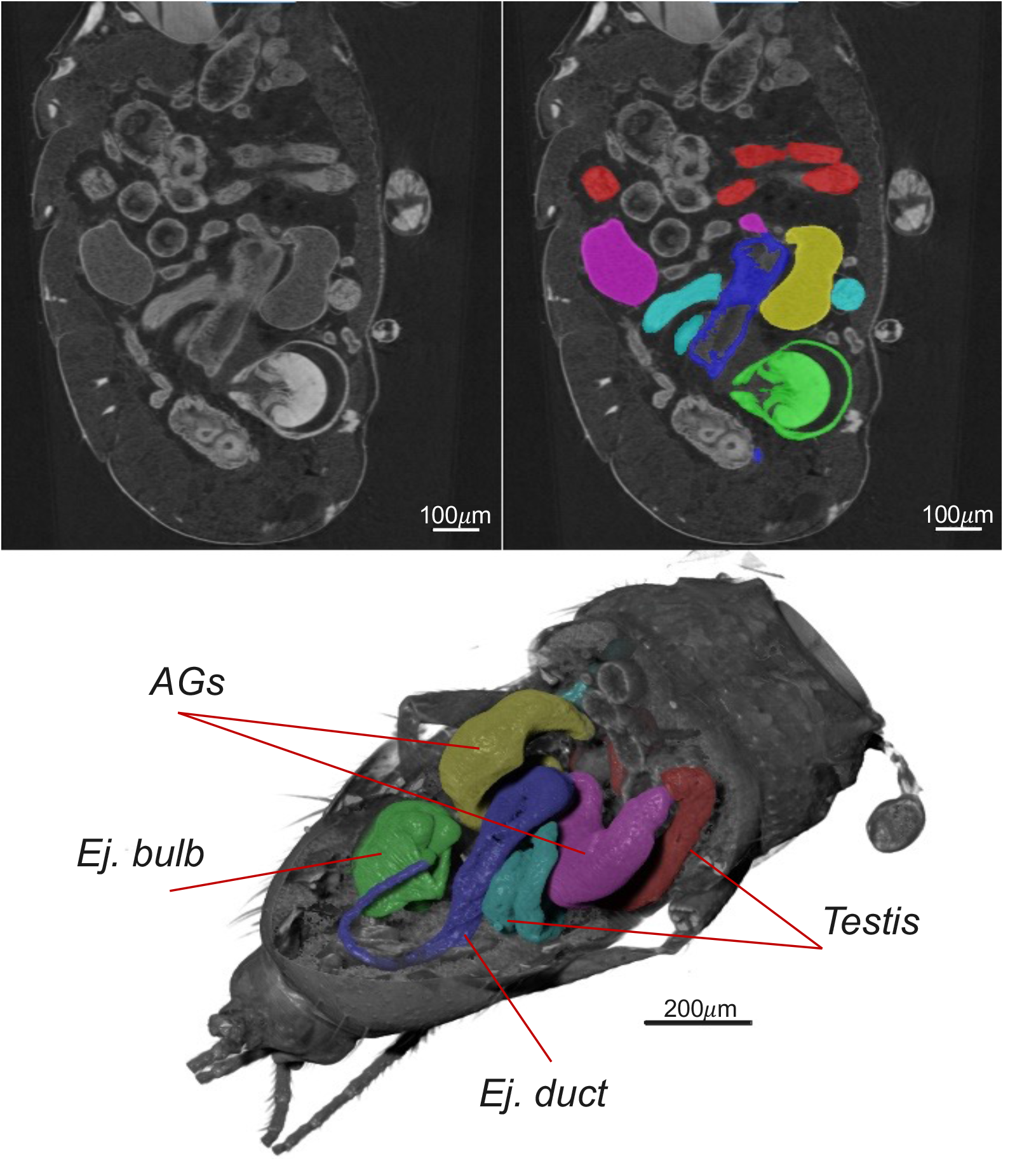
*µ*CT imaging of male reproductive system in *Drosophila*. (a) A slice generated after image reconstruction (b) the same slice as in *a* with segmented organs colored. (c) 3D image reconstruction genereted after segmentation.

**Fig 2.**
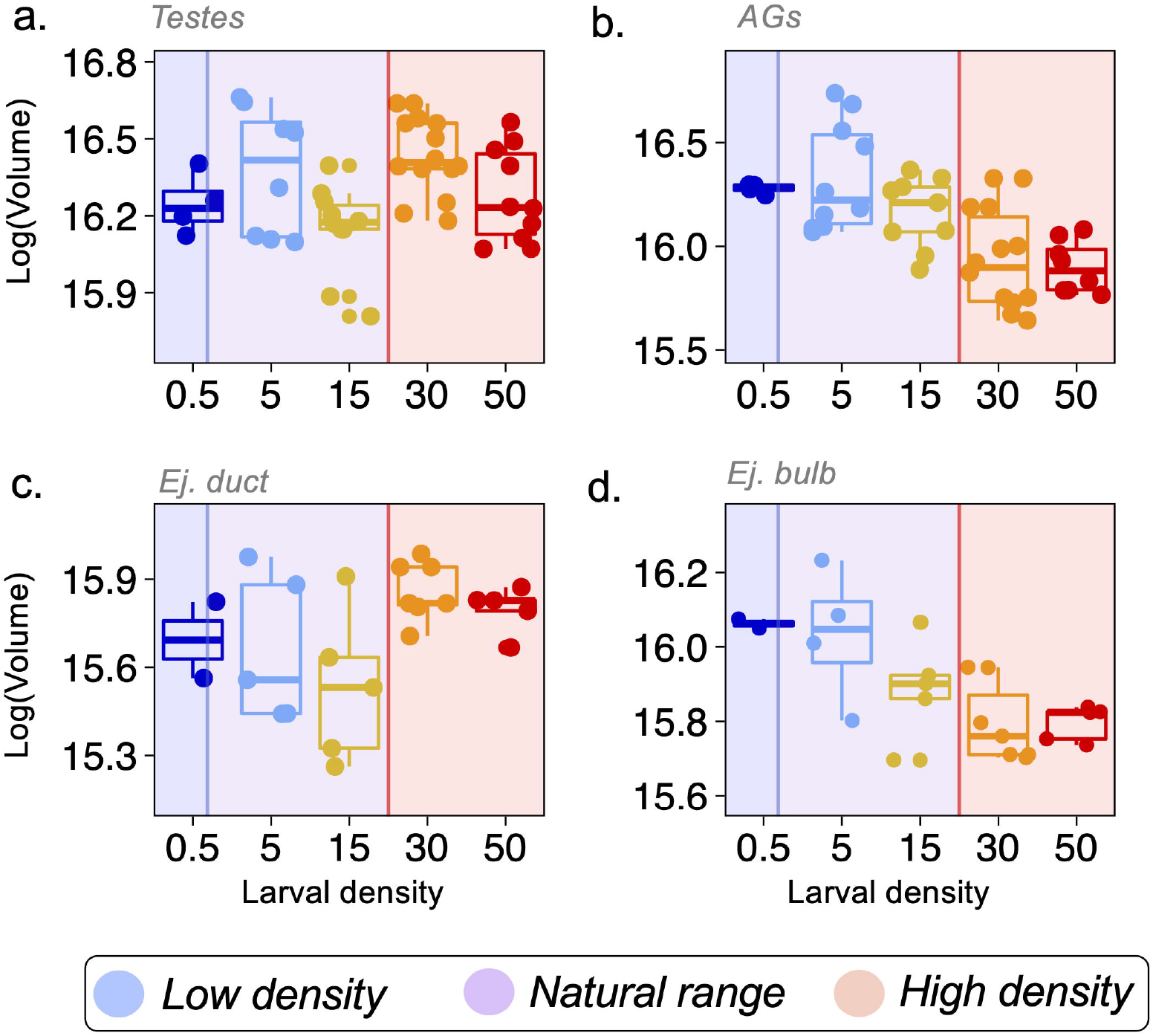
*µ*CT reveals the effect of larval density on male reproductive system. Comparisons across larval density treatments (larvae/g) between the volumes of (a) testes (b) accessory gland (c) ejaculatory duct and (d) ejaculatory bulb. Volume was log-transformed for model fitting. Volume units are given in log(*µ*m^3^). Shaded region indicates the corresponding larval density observed in natural populations, whereby blue = low density, purple = natural range, and red = high density according to (Morimoto and Pietras 2020).

## Discussion

In this paper, we discuss the potential to use micro-CT imaging to empirical studies in evolutionary ecology with a focus on reproductive biology. Micro-CT has proven tremendously useful to studies of insect anatomy, physiology, development, taxonomy, and phylogeny (Carlson 2006; Metscher 2009a; Schambach et al. 2010; Abel et al. 2011; Rawson et al. 2020). However, its use in ecological applications has lagged significantly behind other biological disciplines such as biomedicine, likely due to Micro-CT’s relationship to Computed Tomography (CAT Scans), which were originally designed specifically for medical imaging applications in clinical settings (Gutiérrez et al. 2018). Nonetheless, the technical capabilities of micro-CT allow insects to be imaged in an intact state, preserving the overall spatial architecture of organs in their native orientation. Combined with the isotropic resolution of the technique and powerful segmentation tools, which allow for highly accurate quantitative measures of morphometry (Schoborg et al. 2019; Schoborg 2020), we suggest that micro-CT can be a useful tool for answering ecological questions related to understanding an insect’s response to its environment. In fact, 3D imaging is changing the way in which we understand the anatomy of organs across orders of insects, particularly through recent studies of brain atlases [see (Adden et al. 2020; Rother et al. 2021) and references therein]. Thus, micro-CT is a powerful technique that, combined with ecological experiments, can reveal important responses to changing environments. Recent studies harnessing the power of micro-CT imaging have been transformative to understand plastic and evolutuionary morphological adaptations, particularly in reproductive traits. In *D. melanogaster*, for instance, micro-CT revealed the dynamics of the male-female reproductive structures during mating, showing for example that males’ pierce females’ vagina upon intromission and that mating induces changes in the uterine volume and shape (Mattei et al. 2015). Likewise, in beetles, micro-CT also helped unravel the links between traumatic insemination and female kicking, revealing a temporal offset between the former and the latter behaviours (Dougherty and Simmons 2017), and that females do not respond plastically by thickening their reproductive track in response to high sexual conflict environments (Wyber et al. 2021). Lastly, in lepidopterans, micro-CT has provided important empirical data to support theoretical predictions that female genital teeth morphology, which is associated with a counter-response by females to male reproductive manipulation, evolve in response to the intensity of sexual conflict (McNamara et al. 2019).

From a technical point, it is important to mention that special care during sample preparation is essential to obtain good micro-CT scans (Metscher 2009b). The use of fixation and staining protocols are requisites to study the morphology of soft tissues in the biological samples since the choice of appropriate chemical agents is a simple and effective way to increase the contrast of low-absorbing tissues. It is recommended, depending on the composition and size of the sample, to study the concentration of the staining agent and the time of the process. Recent studies have provided guidelines for the use of different stainings in soft biological matter [see e.g., (Keklikoglou et al. 2019)]. In the entomological context, it is important to know that the insect exoskeleton poses a difficult barrier for penetration by phosphotungstic acid (PTA). On the other hand, iodine presents a faster and well-distributed penetration into the sample containing exoskeleton, although resulting in lower resolution of microstructures. When samples are separated from the entire body such that zones of soft tissues are exposed, then PTA can become a more attractive staining solution for high-resolution micro-CT because of the good penetration and high resolution. For instance, a study has optimised the use of PTA for imaging of bumblebee brain structures, whereby PTA outperformed iodine and other staining solution in terms of resolution of micro-structures, although iodine was faster to perfuse through the samples (Smith et al. 2016). Comparing the advantage and disadvantages of different protocols present in the specialized literature, iodine makes it more suitable for studying whole insect specimens (Metscher 2009b; Du Plessis et al. 2017; Sena et al. 2019). Thus, it is important to balance the pros and cons of each staining method according to the type of sample, costs, time, and the resolution needed for the images.

It is important to mention that, although our sample size was relatively small, it is well within – and in the upper range of – sample sizes in the past literature. Previous studies with more than two treatment groups have used sample sizes ranging from *N* = 2 to *N* = 5 [see e.g., (Mattei et al. 2015; Dougherty and Simmons 2017)]. Other studies had larger sample size per treatment but similar final sample size as the one presented in this study [e.g., (McNamara et al. 2019)]. Few studies have used larger total sample sizes, which are still smaller than 2-fold the sample size of our study (Dougherty et al. 2017; Wyber et al. 2021). In this study, our sample size ranged from *N* = 2 to 7, and thus, within the range used in the published literature. While powerful, micro-CT can still be expensive to run particularly for acquiring high-definition images as the one collected here, which limits our (the authors but also the field) to collect large amounts of data comparable to behavioral assays using insects. Nevertheless, our findings provide important insights into the morphological responses to the intraspecific competition levels in the developmental environment in *Drosophila*, and will stimulate further studies using this technique on the topic.

In this study, to further demonstrate the potential for micro-CT to advance our understanding of evolutionary ecology, we manipulated *D. melanogaster* larval density, based on larval population levels observed in a natural population, and showed a tissue-specific response to larval density in adult males. Previous studies in *D. melanogaster* have found that larval density does not affect sperm size (Amitin and Pitnick 2007) nor the transcription activity of seminal fluid proteins in the accessory gland (McGraw et al. 2007), suggesting that changes in ejaculate quality caused by larval density can be based upon anatomical differences that can affect overall ejaculate composition (e.g., total sperm number, total seminal fluid concentration). However, contrary to our expectations, reproductive organ volume was not always positively associated with larval density, suggesting that, if our assumptions are correct, responses to larval density are more nuanced than previously expected. For instance, while testes volume tended to increase with larval density, accessory glands and ejaculatory bulb volumes tended to decrease with increasing larval density. These differences could underpin a trade-off in ejaculate composition, whereby the ejaculate composition is sperm-biased for males from higher larval densities but balanced and seminal fluid-biased for males from natural range and larval densities, respectively [assuming reproductive organ size or volume reflects ejaculate composition (Gage 1995; Stockley et al. 1997; Stockley and Seal 2001; Lemaitre et al. 2011; Ramm et al. 2015)]. Whether or not the ratio of ejaculate components is dependent upon larval density remains to be investigated. The specific functions of the male ejaculatory bulb and duct have not yet been fully uncovered and thus it is challenging to relate our findings to functional aspects of these organs. Nonetheless, both ejaculatory duct and tract produce important components of the male ejaculate (Heinstra and Thörig 1982; Lung et al. 2001; Bretman et al. 2010) and respond dynamically to mating (and potentially to ecological cues) (Cohen and Wolfner 2018). As discussed above, our study had limited sample size, with sample size per treatment ranging from two to seven. Therefore, it is possible that a larger sample size could result in statistical significance for (some) of the trends observed in the data.

*Drosophila* males are known to modulate ejaculate composition and expenditure based on ecological cues of intraspecific competition in adults [e.g., (Bretman et al. 2010; Hopkins et al. 2019)] in ways that can favour male’s exploitation of rivals’ ejaculate (particularly of costly seminal fluids) under competitive scenarios (Hodgson and Hosken 2006; Sirot et al. 2011). This is in agreement with sexual selection theory which predicts that male ejaculate expenditure should increase with the risk of post-copulatory sexual selection, but decrease with increasing intensity of post-copulatory sexual selection (Parker and Pizzari 2010; Parker et al. 2013). Thus, if males exploit rivals’ seminal fluid, then it is possible that males from high-density environments (where competition is more intense but there is more opportunity for exploitation) (i) have lower cost *per* ejaculate due to the lower production of seminal fluid proteins and (ii) have higher sperm production to enhance efficiency of ejaculate exploitation. Our results suggest testes volume are higher (and accessory glands volume lower) for males from high larval density, which corroborates the idea that under high intraspecific competition levels, male ejaculate should be sperm-biased to enhance ejaculate exploitation. This contradicts the findings that males from high larval density invest relatively more *per* mating opportunity (Wigby et al. 2015). Our data do not allow us to investigate the causes of this contradiction, but it is worth mentioning that the larval densities used in (Wigby et al. 2015) are at least four times higher than the highest density used in this study and that of observed a natural population (Morimoto and Pietras 2020). Such a high larval density could result in higher non-adaptive responses to ejaculate investment per mating opportunity as a response to extreme intraspecific competition levels. Nevertheless, our results are broadly consistent with the wider holometabolous insect literature (Than et al. 2020). For example, males from high larval density developed larger testes in the yellow dung-fly *Scatophaga stercoraria* (Stockley and Seal 2001) and the moth *Plodia interpuctella* (Gage 1995).

## Conclusion

Micro-CT can be an important tool to understand how organisms adapt morphologically to ecological and evolutionary conditions (Dougherty and Simmons 2018). Using micro-CT, we show that larval density leads to a tissue-specific effect on male reproductive organs, which agrees with predictions from sexual selection theory and contributes to our understanding of the anticipatory morphological responses across larval intraspecific competition levels. Future studies across species using micro-CT will reveal whether the responses found in *Drosophila* are consistent across insects, and can help elucidate why some groups [e.g., Coleoptera (Gay et al. 2009)] appear to differ in their response to larval density compared to others (e.g., Lepidoptera and Diptera) (Than et al. 2020).

## Ethics approval

Not applicable for research with invertebrates.

## Consent to participate (include appropriate statements)

Not applicable

## Consent for publication

Not applicable

## Data availability

Raw data will be available as supplementary information upon acceptance of the manuscript.

## Code availability

Not applicable

## Authors’ contributions

JM designed and conducted the experiment, analyzed the data, wrote the first draft, revised the manuscript. RB performed the segmentation of the regions of interest, quantification and image processing. TAS assisted with identification of structures and editing of the manuscript. LPN performed the chemical staining of the samples, micro-CT scannings, data analysis, study design and writing and editing. MVC designed the micro-CT experiment, conducted data analysis and editing. All authors contributed to the revision of the manuscript and agreed on the final version submitted to the journal.

## Acknowledgements

We would like to thank Dr Stuart Wigby for his useful comments in the early version of this manuscript.

